# Easy multiple sequential CRISPR/Cas9 knockouts in cell lines using a Cre/LoxP re-cyclable vector

**DOI:** 10.1101/2021.02.12.431046

**Authors:** Li Dong, Nima Etemadi, David L Vaux

## Abstract

To easily generate cell lines lacking multiple proteins, we inserted Cre/Lox sites flanking the guide RNA, Cas9, and mCherry fluorescent protein coding regions of a CRISPR/Cas9 lentiviral vector. Cells bearing an inducible Cre recombinase construct can be transfected with the CRISPR/Cas9 lentiviral vector, and mCherry positive cells sorted by flow cytometry. Induction of Cre causes deletion of the guide RNA, *Cas9*, and *mCherry* genes, so that mCherry negative cells can be isolated. After confirming successful targeting of the gene, the cells can be re-infected with the same vector bearing a different guide RNA, and mCherry-positive cells sorted once more. In this way, multiple genes can be mutated sequentially using the same vector and selection marker, without persistent expression of the guide RNA or Cas9. We used this system to sequentially mutate two candidate genes, *Bak1* and *Bcl2*, and generated lines that lacked expression of both proteins.

---

The Clustered Regularly Interspaced Short Palindromic Repeats (CRISPR) associated protein 9 (Cas9) system has been widely adopted as a tool for genetic editing in many species ^1–3^. A user-defined guide RNA (gRNA) directs Cas9 to the desired place in the genome where it creates a double-stranded break (DSB), in which DNA repair can result in frameshifts that inactivate the target gene ^4–6^.

Although the CRISPR/Cas9 system is useful for mutation of genes in cell lines, targeting of several genes in the same cell line can be inconvenient due to the requirement for multiple selection markers. Furthermore, the persistence of expression of Cas9 and the guide RNA can sometimes cause off-target mutations ^7–9^, and can prevent re-introduction of the targeted gene.

Cre-LoxP technology has been widely used to conditionally delete DNA segments. Cre recombinase recognizes 34-bp sequences (5-ATAACTTCGTATAatgtatgcTATACGAAGTTAT-3) referred to as loxP sites, and mediates recombination between two loxP sites, thereby deleting the DNA between them ^10–12^.

Combined use of Cre/LoxP and CRISPR/Cas9 technologies has been used to produce conditional gene knock out lines ^13, 14^. However, these methods did not provide a convenient way to generate multiple sequential CRISPR/Cas9 knockouts. Here we describe a new system using both CRISPR/Cas9 and Cre/LoxP to successfully mutate both *Bcl2* and *Bak1* genes in WEHI7 suspension cell lines, in which sgRNAs, Cas9 and mCherry coding regions could be deleted using Cre, so that further genes could subsequently be mutated using the same CRISPR/Cas9 lentivral vector bearing a different guide RNA.

This system uses two lentiviral vectors. We made the LentiCRISPRv2-loxP-sgRNA-Cas9-loxP vector (**Figure 1a**) by modifying the LentiCRISPRv2-mCherry vector (Addgene plasmid # 99154 from Agata Smogorzewska), which expresses a single sgRNA driven by a U6 promoter, with the Cas9 and mCherry genes driven by an EF1a promoter. Cre mediated recombination of the two inserted loxP sites allows deletion of the sgRNA, Cas9, and mCherry coding regions. **Figure 1b** shows the Tet-On 3G *Cre* EGFP lentiviral vector in which expression of Cre can be induced by addition of Doxycycline (Dox).

**Figure 1.**
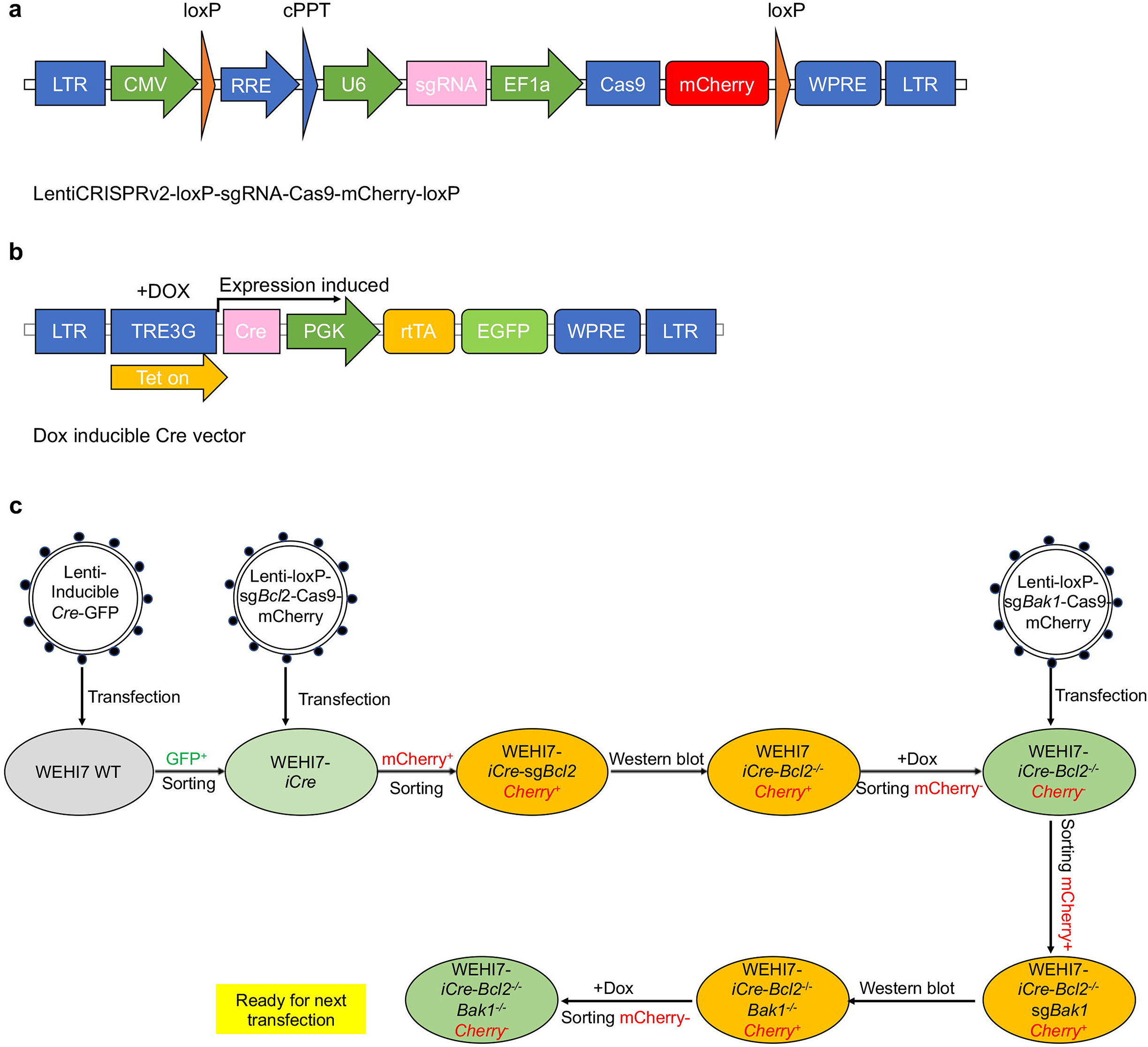
**a.** The LentiCRISPRv2-loxP-sgRNA-Cas9-mCherry-loxP vector. sgRNA (pink) driven by U6 promoters (green arrows) was used to target the corresponding gene. Cas9 and mCherry were driven by an EF1a promoter. Cre mediated recombination of the two loxP sites (orange arrows) can be used to delete the sgRNA, Cas9, and mCherry coding regions. cPPT: central polypurine tract. RRE: reverse response element. LTR: long terminal repeats. WPRE: Woodchuck Hepatitis Virus Posttranscriptional Regulatory Element. **b.** The doxycycline (Dox) inducible Lentiviral Tet-On 3G Cre expression vector. This construct uses the PGK promoter to constitutively express reverse Tet activator (rtTA) and EGFP, and when Dox is added, the Tet-On 3G (TREG3G) promoter to drive expression of Cre. **c.** Schematic of experimental setup. Wild type (WT) WEHI7 cells were transfected with the lentiviral vector encoding EGFP and inducible Cre proteins. Stably transfected cells were isolated by using flow cytometry to detect EGFP fluorescence. The resulting cells were then transfected with the CRISPR/Cas9 lentiviral vector targeting *Bcl2*. Individual mCherry positive cells were sorted by flow cytometry, and the absence of BCL2 was subsequently confirmed by Western blotting of lysates of the expanded clones. Independent clones lacking BCL2 protein were then treated with doxycycline to induce Cre expression, and mCherry negative cells were sorted by flow cytometry for further experiments. Further rounds of CRISPR/Cas9 gene targeting were performed by repeating the process. The *Bcl2* mutant cells were then transduced with the CRISPR/Cas9 lentiviral vector targeting *Bak1*, and sorted for mCherry positive cells, that were subsequently checked for absence of BAK1 by Western blot. Once the targeting mutations have been achieved, the guide RNA, Cas9, and mCherry coding regions can be deleted from the cells by activating expression of Cre recombinasen using Dox. The cells can be re-infected with the same vector bearing yet further guide RNAs, and sorted by flow cytometry once again to detect mCherry-positive cells.

An experimental schema for the generation of *Bcl2^−/−^Bak1^−/−^* clonal lines of WEHI7 thymoma cells is depicted in **Figure 1c**. We first established the WEHI7 *iCre* cell line by transfecting wild type (WT) WEHI7 cells with the Dox inducible lentiviral Tet-On 3G Cre expression vector, and sorted EGFP positive cells using a FACSAria Fusion flow cytometer.

To test the efficacy of this system for generation of multiple sequential CRISPR/Cas9 knockouts in WEHI7 cells, we chose *Bcl2* and *Bak1* because sgRNAs against these genes have been proven to work well in WEHI7 cells ^15^.

First, the Lenti-loxP-*Bcl2* sgRNA-loxP vector was transfected into the WEHI7 *iCre* cell line, and after 48 hrs culture, individual mCherry positive cells were sorted into a 96 well plate by FACS single cell sorting. Once the mCherry positive lines had expanded, the levels of BCL2 protein in the Lenti-*Bcl2* sgRNA-loxP transfected lines was evaluated by Western blot. In 16 lines tested, we obtained 12 independent clones that in which BCL2 protein was undetectable (**Figure 2a**).

**Figure 2.**
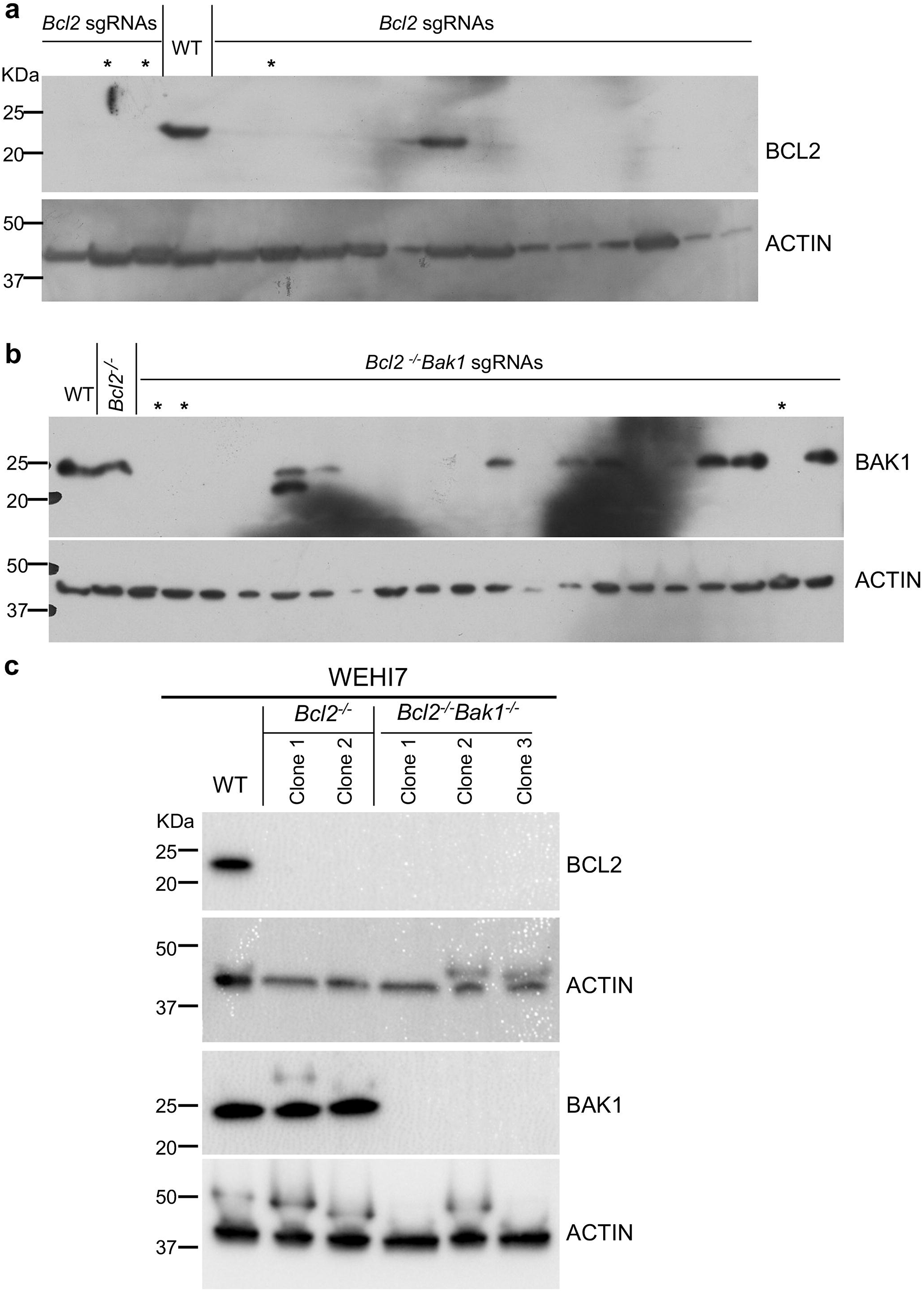
Generation of Bcl2^−/−^Bak1^−/−^ clones by repeated targeting with a CRISPR/Cas9 lentiviral vector that could be recycled using inducible Cre. **a.** WT WEHI7 cells bearing a Dox inducible Cre expression construct were transduced with the lentiviral vector bearing an sgRNA targeting *Bcl2*, as well as genes for Cas9 and mCherry fluorescent proteins. mCherry positive clones were isolated by flow cytometry, and the absence of BCL2 was confirmed by Western blot. BCL2 protein was not detected in 12 of 16 clones examined, and * indicates *Bcl2* deleted clones selected for further experimentation. **b.** Following treatment with Dox to induce Cre expression, *iCre Bcl2*^−/−^ cells that were mCherry negative were sorted by flow cytometry, and were then transduced with the lentiviral vector bearing an sgRNA targeting *Bak1*. mCherry positive clones were sorted by flow cytometry, and the absence of BAK1 was confirmed by Western blot. * indicates *Bcl2* and *Bak1* mutant clones that were selected for further experimentation. **c.** Whole cell lysates from WEHI WT and 2 independent *Bcl-2*^−/−^, and 3 independent *Bcl2*^−/−^*;Bak1*^−/−^ WEHI7 cell clones were analysed by Western blot to detect BCL2 and BAK1 proteins. Aliquots from the same lysates were run on replicate gels and probed for ACTIN to indicate loading.

We then treated the *Bcl2*^−/−^ WEHI7 *iCre* cells with Dox for 48 hrs, and analysed them by flow cytometry. As shown in **Figure 3a**, after 48 hrs of induction of Cre, almost all of the cells became mCherry negative. In one of the WEHI7 i*Cre Bcl2*^−/−^ lines, some mCherry negative cells appeared without the addition of Dox, consistent with some leakiness in the expression of Cre recombinase ^16, 17^.

**Figure 3.**
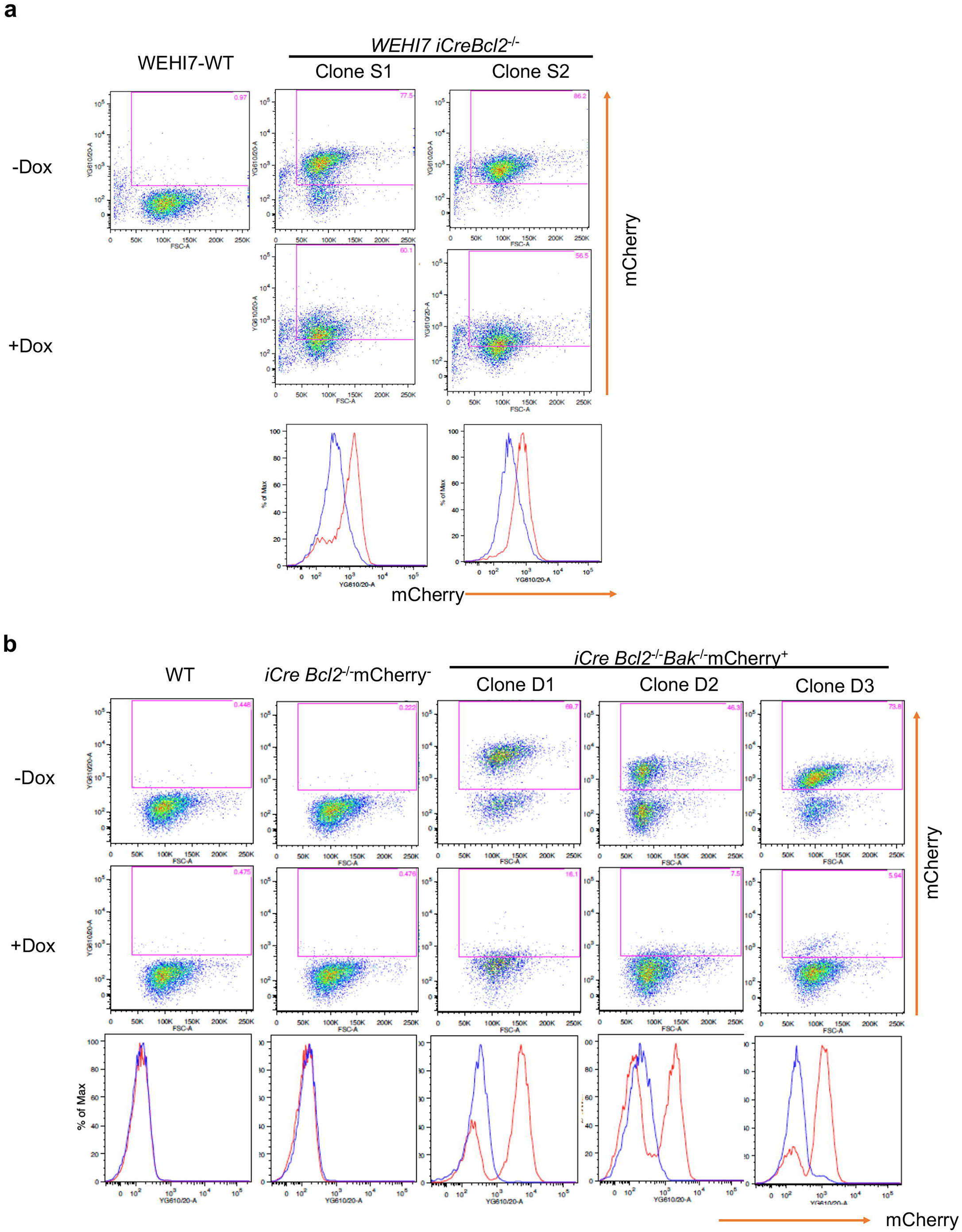
Cre-loxP mediated deletion of mCherry, Cas9 and sgRNA encoding sequences. **a.** Wild type (WT) WEHI7 cells and two independent *Bcl2*^−/−^ WEHI7 cell clones (S1, S2) bearing the inducible Cre (*iCre*) vector in addition to the CRISPR vector were cultured for 2 days in the presence or absence of 1 µg/ml doxycycline (Dox). 2D dot plots show mCherry expression assessed by flow cytometry. Histogram profiles (below) show cells treated with Dox (blue lines) or without Dox treatment (red lines). Treatment with Dox caused all of the cells to lose mCherry fluorescence due to CreLox mediated deletion of the sgRNA, Cas9, and mCherry encoding regions of the vector. **b.** One *Bcl2*^−/−^ clone in which the CRISPR vector had been deleted, but still bore the inducible Cre (*iCre*) vector, and three *Bcl2*^−/−^ *iCre* clones that had been re-infected with a CRISPR vector targeting *Bak1,* were cultured for 2 days in the presence or absence of 1 µg/ml Dox. 2D dot plots show mCherry expression assessed flow cytometry. Although there was some spontaneous loss of mCherry expression in the absence of Dox, presumably due to leaky Cre expression, addition of Dox caused all the cells to become mCherry negative. Histograms (below) compare cells treated with Dox (blue lines) with cells cultured without Dox (red lines).

After 48 hrs treatment with Dox, mCherry negative WEHI7 *iCre Bcl2*^−/−^ cells were sorted by flow cytometry. We then transfected these cells with the Lenti-loxP-*Bak1* sgRNA-loxP vector, and after 2 days culture, we isolated mCherry positive cells by FACS single cell sorting. After expansion of these cells, Western blot analysis revealed that of 20 independent clonal lines tested, 9 lacked detectable BAK1 (in addition to lacking BCL2) (**Figure 2b**). Thus, we successfully generated WEHI7 cell lines with double knock-out of *Bcl2* and *Bak1* genes (**Figure 2c**).

To confirm that mCherry fluorescence could be removed in the WEHI7 *iCre Bcl2*^−/−^ *Bak1*^−/−^ clones upon Cre activation, we treated the cells with Dox for 2 days and examined them by flow cytometry. As anticipated, treatment with Dox caused loss of mCherry fluorescence in all the three independent *iCre Bcl2*^−/−^ *Bak1*^−/−^ clones (**Figure 3b**).

In summary, we present here an easy method for generation of multiple sequential CRISPR/Cas9 knockouts in cell lines. This system allows the same vector to be re-used without the need for additional selection markers. Because the sgRNA, Cas9 and mCherry fluorescent protein coding regions were flanked by the two loxP sites, once the targeting mutation has been achieved, they can be removed by inducing expression of Cre recombinase. This enables the cells to be re-infected with the same vector bearing further sgRNAs, such that those transfected can be sorted by expression of mCherry once again. In addition, this system prevents prolonged expression of Cas9 and the sgRNA, reducing the likelihood of off-target effects, because these sgRNAs and Cas9 coding regions can be removed by activating expression of Cre recombinase.

## Materials & Methods

### Cell culture and reagents

HEK 293T cells were cultured in DMEM (Thermo Fisher Scientific, Cat# 11885-084) supplemented with 8% fetal bovine serum (FBS, Sigma Cat# F9423). WEHI7 cells were cultured in RPMI 1640 medium (Life Technologies) supplemented with 8% fetal bovine serum. Cells were cultured at 37 °C and 10% CO_2_. All cell lines used were routinely tested for mycoplasma contamination.

Doxycycline were purchased from Sigma-Aldrich.

### Plasmid construction and inducible expression

LentiCRISPRv2-mCherry was a gift from Agata Smogorzewska (Addgene plasmid # 99154; http://n2t.net/addgene:99154; RRID: Addgene_99154). In order to insert the two loxP sites, we designed two separate fragments, F1 and F2. F1 contained a loxP site and NotI and KpnI restriction sites. F2 contained a loxP site and BsrGI and SacII restriction sites. The F1 and F2 fragments were synthesised by Integrated DNA Technology (IDT). The fragments were amplified and sub-cloned into a TA vector. The F1 and F2 fragments were digested out of TA vectors using their respective restriction enzymes and then ligated to LentiCRISPRv2-mCherry in two steps. The target sequences of sgRNAs for *Bcl2* and *Bak1* were described previously ^15^. Complementary oligonucleotides were annealed and cloned into the BsmBI sites of the LentiCRISPRv2-loxP-sgRNA-Cas9 plasmid.

For inducible Cre expression, the doxycycline inducible lentiviral vector pFTRE 3G rtTA EGFP/Cre, kindly provided by Prof. John Silke (WEHI), was used to infect cells, and clonal lines were obtained by sorting single EGFP positive cells.

### Preparation of lentivirus

HEK293FT cells were transfected with a lentiviral vector with pVSV-G and pCMV8.2 using Effectene (QIAGEN, Hilden, Germany). Forty-eight hours after transfection, supernatant containing lentivirus was collected for spin-infection as described previously^18^.

### CRISPR/Cas9 gene mutation

For CRISPR/Cas9 gene deletion, parental cells were infected with lentiviral constructs encoding Cas9 and mCherry, and a single guide RNA targeting the desired gene. Successful transfected cells were isolated by sorting mCherry positive cells on a flow cytometer (FACS, Becton Dickinson). Independent single cell clones lacking the targeted protein were confirmed by immunoblotting.

### Flow Cytometry and cell Sorting

For flow cytometric analysis of mCherry, cells were harvested and then resuspended in phosphate-buffered saline (PBS) before being analyzed by a LSRFortessa X-20 flow cytometer (BD Biosciences San Jose, CA). The threshold for detection was set at 10,000 counting events. Data were analysed using FlowJo software (BD Biosciences, San Jose, CA). Wild-type (WT) WEHI7 cells were used to design gate for analysis with FACS.

For FACS sorting, cell suspensions were resuspended in PBS containing 2% FBS and then separated on a FACSAria Fusion flow cytometer (BD Biosciences, San Jose, CA). For single cell sorting, one cell was seeded per well into flat-bottomed 96 well plates (BD Biosciences) containing 100 μL RPMI 1640 medium supplemented with 8% FCS. At day 14, the clones were moved to 24 well plates (BD Biosciences) for further expansion. Confluent clones were split and replicated plates were harvested for western blot analysis.

### Western blot

Total cell lysate was prepared in RIPA buffer (50 mM Tris-HCl, 0.1% SDS, 1% Nonidet P-40, 0.5% deoxycholate and 150 mM NaCl, pH 8.0) complemented with protease inhibitors. Proteins at the same amount were separated by Tris-glycine gels (BioRad) and transferred onto polyvinylidene difluoride membranes. Membranes were then incubated with blocking buffer for 1 h at room temperature, followed by indicated antibodies for 16 h at 4 °C. After probing with primary antibody, membranes were probed with HRP conjugated anti-IgG secondary antibodies and ECL (GE Healthcare Life Sciences). The antibodies used were: ACTIN (AC-15, Sigma #A1978), BAK1 (aa23-38, #B5897 Sigma), BCL2 (#610539, BD Biosciences).

## Acknowledgements

We thank Prof. John Silke for the inducible Cre construct, and Agata Smogorzewska for the LentiCRISPRv2-mCherry vector. This research received funding from NHMRC grants 1113133 and 1135864 to DLV and was made possible through Independent Research Institutes Infrastructure Support Scheme grant 361646 from the NHMRC and a Victorian State Government Operational Infrastructure Support Grant. The plasmids for the Dox-inducible Cre expression vector and the LentiCRISPRv2-loxP-sgRNA-Cas9-mCherry-loxP vector will be made available through Addgene.

## Author Contributions

DLV conceived and designed the project; NE did the vector design and construction; LD performed experiments; DLV and LD analyzed and interpreted the data; DLV and LD wrote the manuscript.

## Conflict of Interest Statement

The authors have declared that no competing interests exist.

## Notes

### Competing Interest Statement

The authors have declared no competing interest.

